# Novel murine model of human astrovirus infection reveals a cardiovascular tropism

**DOI:** 10.1101/2024.10.03.616429

**Authors:** Macee C Owen, Yuefang Zhou, Holly Dudley, Taylor Feehley, Ashley Hahn, Christine C Yokoyama, Margaret L Axelrod, Chieh-Yu Lin, David Wang, Andrew B Janowski

**Affiliations:** Immunology Program, Washington University School of Medicine, St. Louis, Missouri, USA; Department of Pediatrics, Division of Pediatric Infectious Diseases, Washington University School of Medicine, St. Louis, Missouri, USA; Department of Molecular Biology and Microbiology, Case Western Reserve University, Cleveland, Ohio, USA; Xanadu Bio, Gladwyne, Pennsylvania; Department of Internal Medicine, Division of Dermatology, Washington University School of Medicine, St. Louis, Missouri, USA.; Department of Pathology and Immunology, Washington University School of Medicine, St. Louis, Missouri, USA.; Department of Molecular Microbiology, Washington University School of Medicine, St. Louis, Missouri, USA

**Author notes:** Address correspondence to: Andrew Janowski, Washington University School of Medicine, 425 S Euclid Avenue, MSC-8116-16-6, St. Louis, MO 63110 USA. 1-314-454-6050.

## Abstract

Astroviruses are a common cause of gastrointestinal disease in humans and have been recognized to cause fatal cases of encephalitis. A major barrier to the study of human-infecting astroviruses is the lack of an *in vivo* model, as previous attempts failed to identify a suitable host that supports viral replication. We describe a novel murine model of infection using astrovirus VA1/HMO-C (VA1), an astrovirus with high seroprevalence in humans that is a causative agent of encephalitis. VA1 RNA levels peak in heart tissue at day 7 post-inoculation. The cardiotropism was observed in multiple different murine genetic backgrounds evidenced by high VA1 RNA loads in heart tissue of A/J, C57BL/6, C3H/HeJ, Balb/c, and J:ARC mice. Infectious VA1 particles could be recovered from heart tissue 3 and 5 days post-inoculation. Intracellular viral capsid was present in tissue sections based on immunofluorescent staining and viral RNA was detected in cardiac myocytes, endocardium, and endothelial cells based on fluorescent *in situ* hybridization and confocal microscopy. Histologically, we identified inflammatory infiltrates consistent with myocarditis in some mice, with viral RNA co-localizing with the infiltrates. These foci contained CD3+ T cells and CD68+ macrophages. Viral RNA levels increased by > 10-fold in heart tissue or serum samples from Rag1 or Stat1 knockout mice, demonstrating the role of both adaptive and innate immunity in the response to VA1 infection. Based on the *in vivo* tropisms, we also tested cardiac-derived primary cells and determined that VA1 can replicate in human cardiac microvascular and coronary artery endothelial cells, suggesting a novel cardiovascular tropism in human cells. This novel *in vivo* model of a human-infecting astrovirus enables further characterization of viral pathogenesis and reveals a new cardiovascular tropism of astroviruses.

**Author Summary:** Astroviruses typically cause viral diarrhea in humans but can also cause serious infections of the brain. Previously, the methods available to study how these viruses lead to invasive infections were limited. Here, we describe the first system to study human-infecting astroviruses using mice. We demonstrate that mice are susceptible to astrovirus VA1, a strain that commonly infects humans and has been linked to fatal brain infections. The virus infected heart tissue and was associated with inflammation. When mice with impaired immune systems were infected with VA1, they were found to have higher amounts of virus in their hearts and blood. Linking back to human health, we also found that VA1 can infect cells derived from human blood vessels of the heart. This model will enable us to better understand how astroviruses cause disease and how the immune system responds to infection. Our findings also suggest that astroviruses could be linked to cardiovascular diseases, including in humans. In the future, we can develop interventions that will prevent and treat astrovirus infections in humans.

## Introduction

Astroviruses are a diverse family of RNA viruses that are frequently detected from many vertebrate species, including mammals, birds, reptiles, amphibians, and fish (1). Initially discovered in 1975 from an outbreak of gastroenteritis, astroviruses have been primarily detected from stool specimens and wastewater (2,3). The causal link between their presence and causing gastrointestinal (GI) disease was confirmed by oral challenge studies in humans and other mammals (4–8). Further epidemiological studies have established astroviruses as a common etiological agent of gastroenteritis with some estimates nearing 6 million new human cases each year (9).

Since 2010, astroviruses have received additional attention due to the recognition of their capacity to cause central nervous system (CNS) diseases. In humans, 15 cases have been reported with a mortality rate of >50% (1,10–12). Similar discoveries have been reported in other mammals infected by other astrovirus genotypes, including in cattle, pigs, minks, alpaca, muskox and sheep (13–19). Currently, astroviruses are not part of routine clinical testing, so the disease burden caused by astroviruses in neurological diseases in humans and other mammals is poorly understood.

In addition to the newly described neurotropism, there is increased recognition that astroviruses have additional tissue tropisms (1). Avian astroviruses have been linked to fatal liver and kidney diseases that are of significance to the poultry industry (20–22). In humans, astroviruses have been implicated as a possible cause of hepatitis and increased risk of immune thrombocytopenia (23,24). Astroviruses have also been detected from the respiratory tract; however, further data is needed to clarify whether there is a causal relationship between astrovirus infection and respiratory diseases (14,25–34). Other potential tissue tropisms have been described, including detection of viral nucleic acid from heart tissue in ducks, chickens, geese, pigs, and humans (35–39). However, it is unclear to what extent these findings are incidental findings versus if they reflect *bona fide* cardiovascular infections.

Three clades of astroviruses are recognized to cause disease in humans; the classic human astrovirus (HAstV), MLB, and VA clades (3). Most humans have been exposed to these clades and develop neutralizing antibodies, as >50% of adults have a serological response to at least one species within each clade (40–46). All three clades are frequently detected from human stool samples from cases of diarrhea or gastroenteritis (3). In addition, they all have been associated with meningoencephalitis in humans, with astrovirus VA1 (VA1) being the most frequently detected strain (10–12,25,29,38,47–53).

Despite the significant role of astroviruses in human health, there are only limited experimental systems to study the pathogenesis of disease, including the lack of any *in vivo* model of infection for a human-infecting astrovirus. One group described their attempts to use human-infecting astrovirus strains to inoculate mice, but there was no clear evidence of viral replication or virus-induced disease in the mice (54). Other models using non-human infecting astroviruses have provided important insights into viral pathogenicity. A turkey astrovirus strain causes diarrhea in a turkey poult model, with detection of viral RNA in many tissues including the liver, spleen, kidneys, bone marrow, and plasma (55,56). Replication was hypothesized to be restricted to the GI tract based on detection of intracellular RNA from *in situ* hybridization assays (55,56). Murine astrovirus strains have also been detected from mice and used as an experimental model system (54,57–60). However, the murine astrovirus model has limitations. Most laboratory and commercially available mice are infected with murine astrovirus, and viral infection does not cause any apparent disease (54,57,59).

Given the limitations of current animal models, we sought to develop a murine model of infection using a human-infecting astrovirus. We used a previously propagated VA1 genotype for inoculation into mice(61,62). Unexpectedly, we identified viral RNA, viral capsid, and isolated infectious viral particles from heart tissue that was associated with inflammatory infiltrates. This novel model of human astrovirus infection overcomes a major limitation in the study of astroviruses and enables further study of pathogenesis of infection. Furthermore, it demonstrates a previously uncharacterized cardiovascular tropism for astroviruses.

## Materials and Methods

### Animals

A/J (strain #000646), C57BL/6 (strain #000664), C3H/HeJ (strain #000659), Balb/c (strain #000651), and J:ARC(S) (strain #034608), B6.129S7-Rag1^tm1Mom^/J (strain #002216), and B6.129S(Cg)-Stat1^tm1Dlv^/J (strain #012606) mice were obtained from the Jackson Laboratory. All mice were maintained in a specific-pathogen free facility following institutional guidelines and with protocols approved by the AAALAC-accredited Animal Studies Committee at Washington University in St Louis. All animals were maintained on 12-hour light cycles and housed at 21°C and 50% humidity. Experiments were performed with mice at 5-6 weeks of age and were carried out utilizing BSL2 conditions. Mice were given access to food and water *ad libitum*.

### Cell culture

All cell lines and infection steps were carried out at 37°C with 5% CO2. The undifferentiated human intestinal epithelial cell line (Caco-2) was cultured in growth medium consisting of Dulbecco’s modified Eagle medium (DMEM) with L-glutamine supplemented with 10% fetal bovine serum (FBS; Gibco) and 1% of 10,000 units/mL of penicillin and streptomycin (Gibco). Primary cell lines were purchased from Sciencell, including primary human cardiac microvascular endothelial cells (CMEC), human coronary artery endothelial cells (HCAEC), human umbilical vein endothelial cells (HUVEC), human hepatic sinusoidal endothelial cells (HHSEC), human cardiac myocytes, and C57BL/6 murine cardiac myocytes. C57BL/6 Mouse primary cardiac microvascular endothelial cells were purchased from Cell Biologics. All primary cells were cultured using the media provided by the vendor, including cardiac myocyte medium- serum free for cardiac myocytes (Sciencell), and endothelial cell medium (Sciencell and Cell Biologics). Infection of cells was performed as described previously using a multiplicity of infection of 3 for all other cell lines (62). Cell fractions were collected at reported timepoints post-inoculation in TRIzol for RNA extraction.

### Viral stock

We used a 0.2 µM sterile-filtered VA1 viral stock that was passaged in Caco-2 cells (C-P8). The stock was generated using the same methods as previously described (62). The stock did not have any mutations relative to a previously described stock (C-P7).

### Murine time course experiment of VA1 inoculation

A/J mice were inoculated intraperitoneally (IP) with 1.5x10^7^ focus forming units (FFU) of C-P8 in 200 µL of DMEM. Mock infected mice were IP inoculated with 200 µL of cell-lysate in DMEM containing no virus. At least three independent infections of mice were performed, typically in groups of 3-5 mice per experiment. Mice of both sexes in similar ratios were tested for all experiments. Mice were monitored daily for changes in weight and clinical signs of infection. Mice were euthanized on days 3, 5, 7, 14, or 21 post-inoculation (p.i.), and tissues were harvested.

### VA1 inoculation in mice of different genetic backgrounds and routes

C57BL/6, Balb/c, C3H/HeJ, J:ARC(S) (Swiss outbred), B6.129S7-Rag1^tm1Mom^/J, and B6.129S(Cg)-Stat1^tm1Dlv^/J were inoculated IP with 1.5x10^7^ FFU of C-P8 in 200 µL of DMEM as previously performed for A/J mice. Inoculations were performed in three independent experiments of 3 mice per experiment. Mice were euthanized and tissues were harvested at day 7 p.i. as described in below methods.

Mice were also infected by different routes. For the *per os* route (PO), 100 µL was slowly pipetted into the mouths of the mice. The mice were allowed to lick and swallow the liquid until the full volume was delivered. For oral gastric lavage (OG), a lavage needle was inserted into the oropharynx of the mice and passed until it entered the stomach. A total of 200 µL was then administered. For intracranial inoculations (IC), mice were anesthetized with a ketamine/xylazine cocktail (100 mg/kg for ketamine, 10 mg/kg for xylazine). Once the mice were sufficiently sedated, 20 µL (1.5x10^6^ FFU) of inoculum was delivered to the mouse central nervous system by insertion of an insulin needle into the cranium. After insertion, mice were monitored for bleeding, neurological deficits, and recovery from the anesthesia cocktail.

### Tissue collection

Mice were euthanized with CO2 and subsequent cervical dislocation.

Blood was obtained from the inferior vena cava via venipuncture. Apexes of hearts were collected for RNA isolation, and the remainder of the heart was transversely sectioned and formalin-fixed for subsequent histopathological analysis. A small section of the left lobe of the liver was excised for RNA isolation, and the entire median lobe of liver was removed and formalin-fixed for histopathological analysis. In addition to the heart and liver, small tissue sections were collected from the following organs: brain, brainstem, lung, kidney, spleen, ileum, colon, mesenteric lymph node, and skeletal muscle. A stool sample was also obtained from the intestinal tract. All tissue samples were weighed and stored in phosphate-buffered saline (PBS) at -80°C. Blood was centrifuged at 6000 RPM for 3 minutes in blood collection tubes (BD), and serum was aliquoted into separate tubes for storage at -80°C.

### RNA isolation

Tissue samples were bead homogenized using a BeadBlaster (BioSpec) in 1 mL of PBS and then centrifuged. RNA was isolated from 100 µL of sample supernatant using TRIzol (ThermoFisher) in the Direct-zol 96 kit (Zymo Research) and stored at -80°C.

### qRT-PCR

A previously published (61) quantitative reverse transcription-PCR (qRT-PCR) was performed to quantify viral RNA from tissue. Isolated RNA was combined with primers, probe, and TaqMan Fast Virus 1-Step master mix (Applied Biosystems) and analyzed on a ViiA 7 or QuantStudio 3 Real Time PCR systems (Applied Biosystems). Viral copy numbers were calculated and normalized to tissue weight by dividing copy number by milligrams of organ weight.

### Focus forming assay

Focus forming assays (FFA) to quantify VA1 titers were performed as previously described (41). Homogenized tissue suspended in PBS was serially diluted for inoculation into Caco-2 cells. The cells were then fixed and stained with the infectious titer quantified.

### Histology

Hearts and livers were collected and fixed in 10% neutral buffered formalin for 48-72 hours. The tissues were then placed in histology cassettes and dehydrated through a series of 70%, 90%, 100% ethyl alcohol, and two rounds of xylenes. Tissues were then paraffin- embedded and sectioned at 4 µM thickness. Tissues were stained with hematoxylin and eosin, examined for any pathological abnormality via light microscopy at 20X magnification, and images were obtained using a Zeiss Cell Observer inverted microscope. Images were edited using the Zen 2.3 lite application (Carl Zeiss Microscopy). Tissue selections were blinded and then independently reviewed for the presence of foci of inflammatory cells by two pathologists. The scoring system for the foci was as follows: 0 = no foci present in the section, 1 = one or two foci present in the section, 2 = three or more inflammatory foci present in the section.

### Immunofluorescent Microscopy

Seven days post-inoculation, mice were anesthetized with ketamine (100 mg/kg) and xylazine (10 mg/kg) before being perfused first with PBS to rinse out the blood and then fixed with 4% paraformaldehyde in PBS (Sigma). The hearts were dissected and post-fixed by immersing in 4% PFA for an additional 3 hours before being cryo- protected by incubation in 15% and 30% sucrose in PBS. The samples were then embedded in Tissue-Tek OCT compound (Electron Microscopy Sciences) and longitudinal sections cut using a cryostat (Cryotome; Leica). Sections were initially washed with PBS, permeabilized with 0.1% Triton-X in PBS, blocked with 5% goat serum/PBS for one hour at room temperature, then incubated with primary antibodies to mouse anti-VA1 capsid (Mab2A2; 1:2000) (41,63), rat anti-CD45 (1:20, BD Pharmingen, #550539), FITC rat anti-CD3 (1:50, Invitrogen, #11-0032-82), or Alexa Fluor 488 rat anti-CD68 (1:200; Biolegend #137011) overnight at 4°C. For anti-VA1 capsid staining, the tissue was first blocked using Mouse on Mouse immunodetection kit (Vector Laboratories), using 2 drops in 1.5 mL of blocking buffer. The sections were incubated with Alexa Flour 488 goat anti-mouse secondary antibody (Invitrogen) at a dilution of 1:1000 for 1 hour at room temperature. For CD45 staining, sections were further incubated with Alexa Fluor 488 goat anti-rat secondary antibody (Invitrogen) at a dilution of 1:1000 for 1 hour at room temperature. Sections were counterstained with 4,6-Diamidino-2-phenylindole (DAPI, Sigma) and imaged at 20-40x using a Zeiss Cell Observer inverted microscope. Images were edited using the Zen 2.3 lite application (Carl Zeiss Microscopy). Four mice for each group were analyzed. A total of five representative images of each heart were obtained and fluorescent cells counted using ImageJ (64). The number of positive cells were normalized to per 1,000 of DAPI positive nuclei.

### Fluorescent *in situ* hybridization (FISH)

FlSH was performed on murine formalin-fixed-paraffin-embedded tissues using RNAscope Multiplex Fluorescent Reagent Kit v2 assay (Advanced Cell Diagnostics; ACDBio) based on the manufacturer’s instructions. As a positive control, VA1-infected or mock-infected Caco-2 cells were collected and suspended in 1% melted agarose. The block was allowed to cool and solidify prior to embedding in paraffin. After sectioning, tissue sections were deparaffinized in xylenes and then dehydrated in 100% ethyl alcohol. Tissues were pretreated with 3% hydrogen peroxide and antigen retrieval was performed in a pressure cooker at 100°C for 15 minutes using RNAscope Target Retrieval Agent diluted to 1X in deionized water. Tissues were treated with RNAscope Protease Plus at 40°C for 30 minutes in an ACDBio RNAscope oven, using the recommended humidity control tray. Targeted RNA sequences were then hybridized using the following target probes obtained from ACDBio: RNAscope Probe V- Astrovirus-VA1-ORF2-C1 (854601-C1), RNAscope Probe-Mm-Vwf-C3 (858851-C3), and RNAscope Probe-Mm-Ryr2-C2 (479981-C2). Positive (RNAscope 3-plex Positive Control Probe-Mm) and negative (RNAscope 3-plex Negative Control Probe-Mm) control probes were utilized to validate the assay. All probes were diluted per manufacturer instructions in Probe Diluent (ACDBio). After washing with RNAscope Wash Buffer diluted to 1X in deionized water, amplification steps were then performed using the RNAscope Multiplex FL v2 AMP1, 2 and 3 per manufacturer’s instructions. Fluorophores obtained from Akoya Biosciences (Opal 520: FP1487001KT, Opal 570: FP1488001KT, Opal 650: FP1497001KT) were reconstituted in DMSO and were diluted at 1:1500 in RNAscope TSA buffer on the same day as intended use. After amplification steps, Opal 520, 570 and 650 were applied to tissues incubated with C1, C2 and C3 probes, respectively. Nuclei were stained with DAPI, and tissues were mounted with Prolong Gold Antifade mounting solution (Thermofisher Scientific). Slides were stored at 4°C, and tissues were visualized using a Zeiss LSM880 confocal laser scanning microscope at 20-63x magnification. Images were cropped and labeled using the Zen 2.3 lite application. Cellular outlines of interest were created based on the staining from the channels and outlined using Photoshop (Adobe). We identified infected cell types based on cells that contained both FISH signal from the VA1 probe and the cell marker probe in the same plane, but subcellular colocalization of the VA1 and cell marker probes was not required as the host and viral RNA can segregate into different locations within a cell (65,66).

### Statistical Analysis

Prism 10.2 (GraphPad) was used for data analysis and graphing.

Comparisons of mouse weights were performed using Mixed effects models. The number of CD68 positive cells from infected hearts were compared to mock infected hearts using a nested T-test within a mixed-effects model. We confirmed there was no statistical difference between mice within the mock or VA1-infected groups. To compare viral RNA loads from immunodeficient mice, Kruskal Wallis testing was used with post-hoc testing to identify significant pairwise comparisons. Adjusted P values ≤ 0.05 were considered significant.

## Results

### VA1 RNA is detectable in murine heart tissue up to 21 days post-inoculation

We first determined whether VA1 could infect and cause disease in wildtype (WT) mice. Five-week-old A/J mice were inoculated by intraperitoneal injection with 1.5x10^7^ focus forming units (FFU) of VA1 or mock infected. Mice were monitored for development of symptoms, weighed daily, and were sacrificed on different days up to 21 days post inoculation (p.i.). No deaths occurred and the mice did not display any clinical signs of illness (hunched posture, lethargy, ruffled fur, decreased activity, seizures, paralysis). Mock-infected mice had a median weight gain of 15% during the observation period. In aggregate, VA1 infected mice also gained weight at a similar rate over 21 days compared to mock inoculated mice (median gain 13%, mixed effects model F[1,97]= 1.84, P= 0.18; Figure 1A). When comparing the weight changes by sex, we noted a significant difference (Figure 1A). Male mice had lower weight gains relative to mock-inoculated mice (Mixed effects model F[1,49]= 6.5, P= 0.014), while there was no difference in female mice (Mixed effects model F[1,46]= 0.42, P= 0.53). 

**Fig 1.**
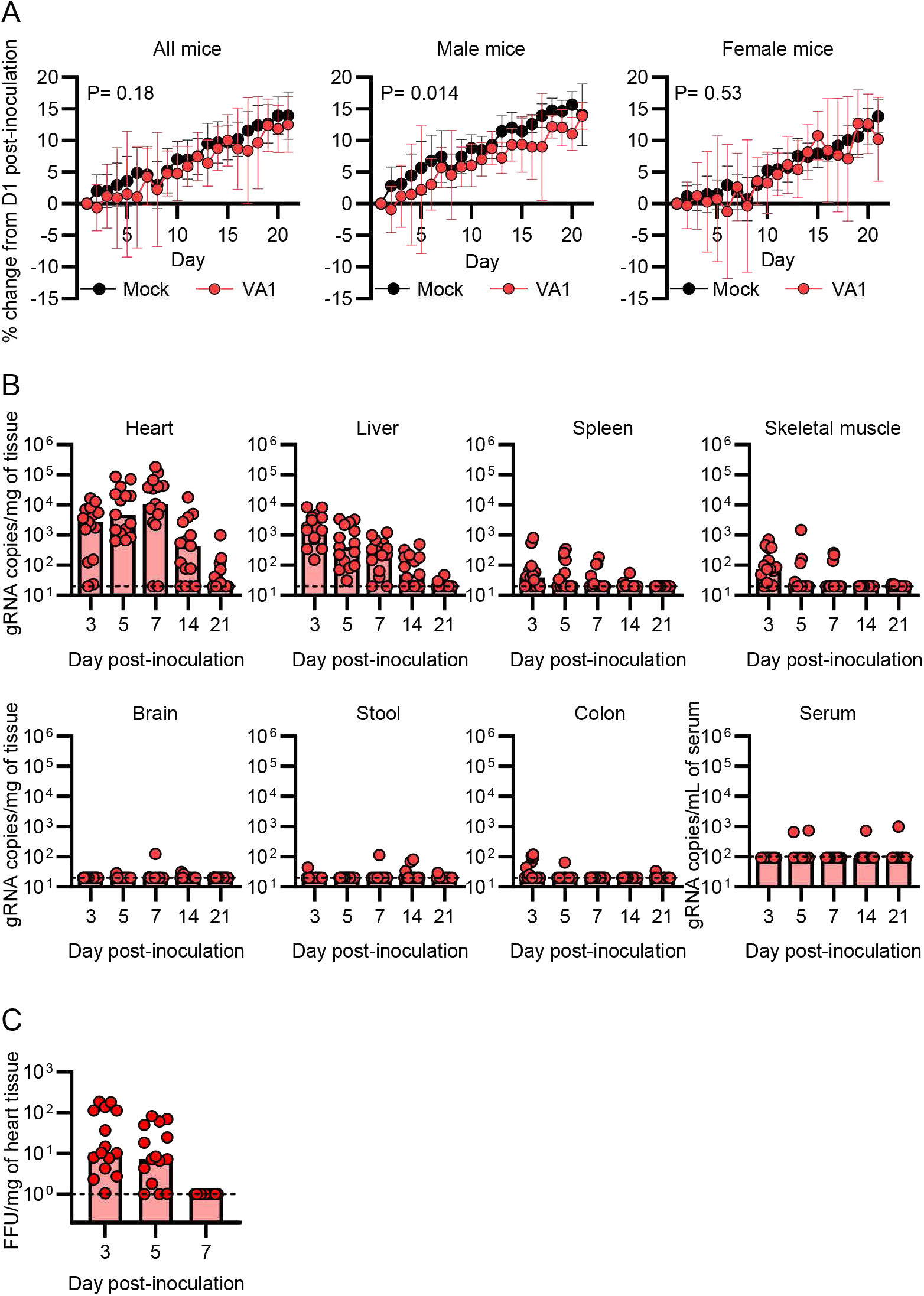
Inoculation of VA1 in wildtype A/J mice results in accumulation of viral RNA in heart tissue. Wildtype A/J mice were intraperitoneally inoculated with VA1. (A) Male but not female mice had reduced weight gain following inoculation with VA1 compared to sex-matched mock-infected mice. (B) Over the course of 21 days post-inoculation, the viral RNA load was measured from tissues. The highest viral RNA loads were in the heart, peaking 7 days post-inoculation but remained detectable in some mice at day 21. Viral RNA was also detected from the liver, spleen, and skeletal muscle while other tissues had sporadic detection of VA1 RNA. Dashed line indicates the limit of detection. (C) Viral infectious titers from heart homogenates collected from VA1- inoculated mice were measured by a focus forming assay. Infectious virus could be detected three and five days post-inoculation, while it was undetectable on day seven. Dashed line represents the limit of detection.

We next determined the kinetics of VA1 RNA over time, and whether inoculation resulted in viral persistence in any particular tissues. Tissues were harvested from mice sacrificed on days 3, 5, 7, 14, and 21 p.i. and viral RNA was measured from the brain, brainstem, heart, lung, liver, kidney, spleen, ileum, descending colon, skeletal muscle, mesenteric lymph node, stool and serum. Remarkably, the highest viral titer was detected in the heart, with >10^3^ gRNA copies per milligram of tissue (copies/mg) at day 3, peaking at 10^5^ copies/mg by day 7, and some mice remained RT-PCR positive at 21 days p.i. (Figure 1B). This remained consistent amongst all experiments with both sexes, with only a small subset of 1-2 of mice out of 15 testing negative from heart tissue at each timepoint in the first seven days after inoculation. Several other organs had consistently detectable virus, including the liver, spleen, and skeletal muscle, including mice that were negative from heart tissue (Figure 1B). In particular, the livers had viral titers of ∼10^3^ copies/mg on day 3 and decreased to near or below the limit of detection by day 21. In other tissues, VA1 RNA was infrequently detected and at low quantities (Figure 1B and S1), including the GI tract, serum, and central nervous system.

We also determined whether infectious virus could be recovered from heart tissue following IP injection. Infectious titers of up to 10^2^ FFU/mg of tissue were detected from heart tissue on day three and five p.i., but no infectious virus was detectable at day 7 (Figure 1C). These results demonstrate that infectious particles are present and recoverable from heart tissue after IP inoculation into a different body site.

### VA1 is tropic to heart tissue in multiple mouse strains

For some viruses, such as influenza and ebolaviruses, different mouse strains can have different capacities to support viral infection (67,68). We used the A/J mouse genetic background to initially characterize the viral kinetics, which is also commonly used to model myocarditis (69). We tested other strains of mice to determine if VA1 tropisms and RNA loads were dependent on genetic background. We inoculated VA1 in three commonly studied inbred strains, including C57BL/6, Balb/c, and C3H/HeJ, and a Swiss outbred strain, J:ARC. All strains demonstrated similar viral loads in the heart tissue compared to A/J mice (Figure 1B and 2A). Like the A/J mice, the other mouse strains also had detectable viral RNA in liver tissue, while the virus was rarely detected from brain and serum (Figure 2A). No deaths occurred and there were no signs of illness. These results further demonstrate VA1’s cardiotropism in multiple mouse strains.

**Fig 2.**
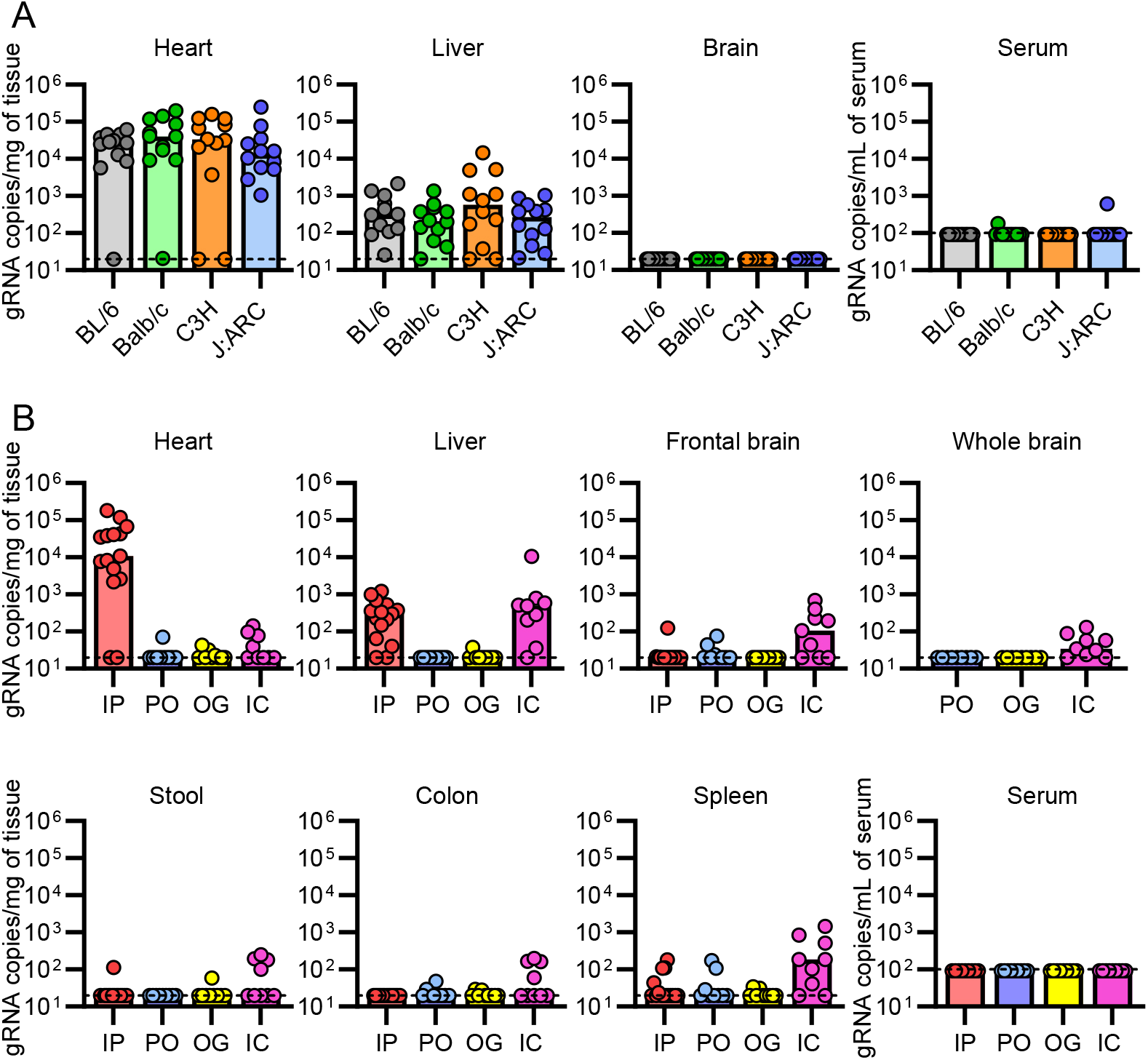
VA1 is cardiotropic in other mouse strains while inoculation by other routes does not result in significant cardiac infection. Commonly studied mouse strains (C57BL/6 [BL/6], Balb/c, C3H/HeJ, and J:ARC Swiss Outbred) were intraperitoneally inoculated with VA1 and viral load was measured by qRT-PCR seven days post-inoculation (A) Like A/J mice, other strains of mice have a significant viral load in heart and liver tissue, while there was only sporadic detection in serum and brain tissue. (B) Different routes of inoculation were tested in wildtype A/J mice, including intraperitoneal (IP; data from Figure 1B, per os (PO), oral gavage (OG), and intracranial (IC) inoculation. Viral RNA loads were measured seven days after inoculation. PO and OG inoculation did not result in significant infection of mice, including in heart tissue compared to IP inoculation. IC inoculation resulted in detection of low quantities of viral RNA from brain tissue, including whole or frontal brain tissue, but was several logs lower than what can be detected in heart tissue from IP inoculation. Viral RNA was also detected from the liver and spleen, while low quantities were detected from the heart. For all graphs, the dashed line is the limit of detection.

### Inoculation of mice by different routes does not result in similar cardiac viral RNA loads compared to intraperitoneal inoculation

We also assessed the impact of other routes of inoculation on tropism of VA1. The fecal-oral route is one of the primary routes of transmission by astroviruses (4–8). Recently, the salivary tract has also been reported as a key site of viral replication for astroviruses and other enteric viruses (*per os* route) (70). We tested these routes separately by inoculating A/J mice by oral gastric lavage (OG), and by pipetting inoculum into the mouths of the mice to test the *per os* (PO) route. Virus was undetectable in most mice on day 7, with only a small subset having a very low viral load in any tissue, including the heart, and no evidence of viremia (Figure 2B). The virus was undetectable in most mice from the GI tract and stool using either PO or OG routes (Figure 2B).

Given the role of VA1 in causing encephalitis, we also tested the intracranial (IC) route of inoculation. The murine blood-brain barrier may prevent dissemination of the virus to the central nervous system, and the IC route would bypass this potential limitation. Inoculation of mice by IC resulted in detectable but low copy numbers from frontal brain tissue and whole brain (Figure 2B). However, these copy numbers were several logs lower than what we observed in heart tissue by IP inoculation, despite direct injection into the brain (Figure 2B) suggesting that replication, if it did occur, is limited. While the virus was not detectable in serum on day 7, we did detect the virus in the liver and spleen (Figure 2B). These results are consistent with dissemination of the virus out of the central nervous system and transportation to these locations, likely mediated by lymphatics, blood, or circulating immune surveillance cells. While this could reflect some limited viral replication, it likely represents immune clearance as the liver and spleen are important organs that mediate this function (71).

### Intracellular VA1 capsid and RNA are detected in heart tissue, with RNA localizing to cardiomyocytes and endocardial cells

Based on the high abundance of viral RNA in heart tissue, we next determined if we could detect VA1 intracellularly. Detection of intracellular VA1 would further validate that VA1 has entered cells and is not transiently passaging through the tissue or extracellularly attached to cells. We first used an antibody to the VA1 capsid in immunofluorescence. We found that VA1 infected mice had areas with positive staining with punctate signal but unclear cellular borders (Figure 3A). These regions appeared to have many nuclei present, as indicated by the DAPI staining (Figure 3A). This result further validates that VA1 infects cells of the murine heart and results in translation of the viral capsid.

**Fig 3.**
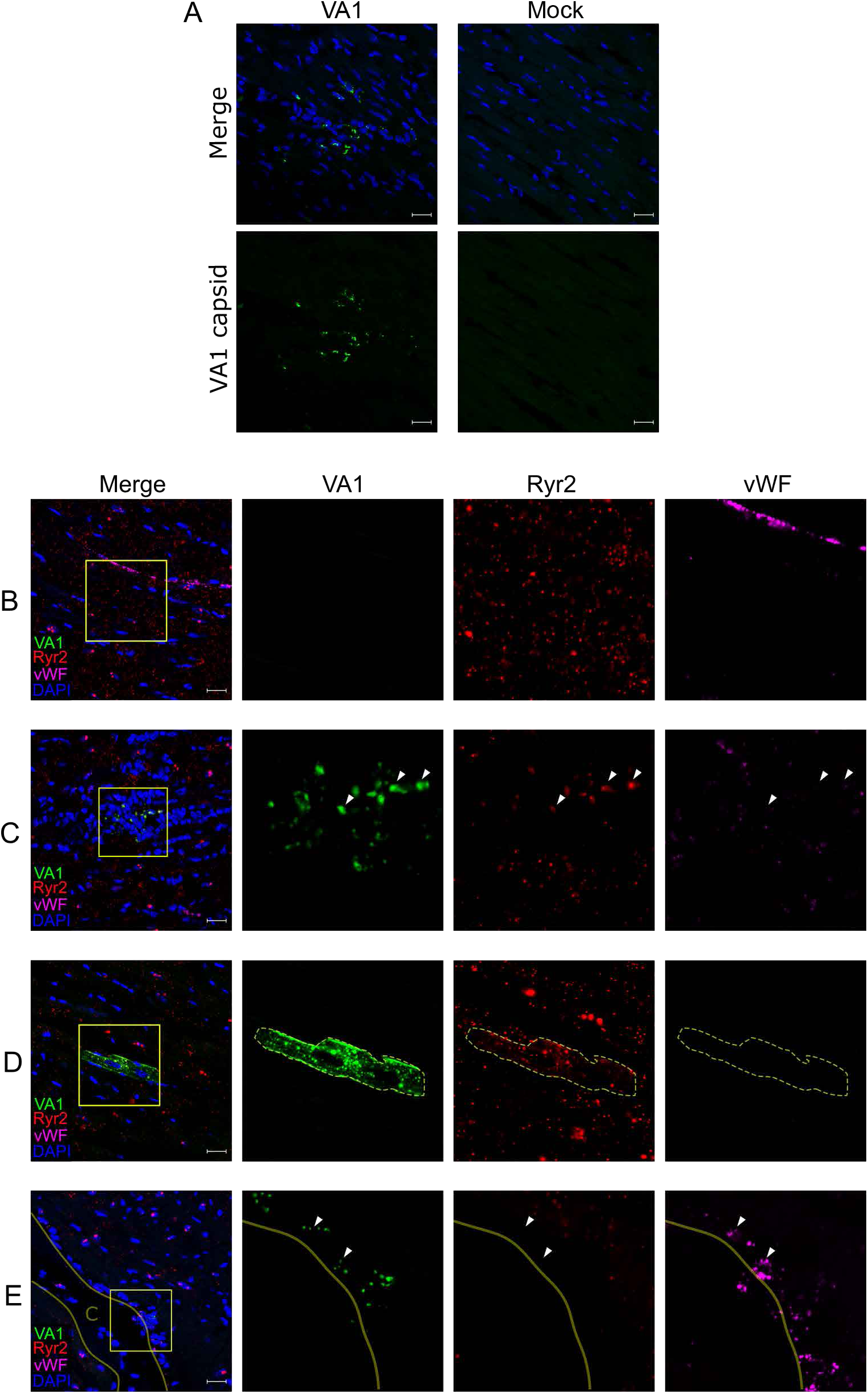
VA1 RNA intracellularly localizes to cardiac myocytes and endocardial cells in wildtype A/J mice. (A) Immunofluorescence staining of VA1- or mock-inoculated mouse hearts. Tissues were processed for immunofluorescence and stained with an antibody to the VA1 capsid (green) and counterstained with DAPI. VA1 infected mice had positive punctate signal in regions with increased nuclei. No staining was observed in mock inoculated mice. (B-E) A fluorescent *in situ* hybridization (FISH) assay using probes specific to VA1 open reading frame 2 (ORF2; green) was developed and imaged by confocal microscopy. After 7 days post-inoculation, heart tissue was obtained and stained with probes to VA1 and to host markers for cardiac cell types including cardiomyocytes (Ryr2 probe; red), and endothelial cells (vWF probe; magenta). Merged fluorescence images are shown with counterstaining of nuclei (performed with DAPI). Boxes demonstrate areas of interest that were further magnified to demonstrate the signal from individual fluorescent channels. Outlines indicate cellular borders of cells containing VA1 RNA. White triangles highlight further areas of staining by the VA1 and host marker probes (B) No staining for VA1 was detected in mock-infected mouse hearts. (C-E) VA1-infected hearts stain positive for detectable VA1 colocalizing with (C) within foci of dense cellular infiltrates and colocalizing with Ryr2, (D) cardiomyocytes expressing Ryr2 without cellular infiltrates, and (E) within a region of endocardial cells expressing vWF. The solid line and the label C denote the open space of the heart chamber. Scale bars represent 20 µm.

To complement the positive intracellular capsid staining results, we developed a VA1 fluorescent *in situ* hybridization (FISH) assay with confocal microscopy to detect intracellular VA1 RNA and to be used in multiplex with other probes to cellular markers. VA1 specific probes were synthesized to the region of open reading frame 2 (ORF2). This region encodes virus capsid precursor protein and is predicted to be present in both genomic and subgenomic RNA strands (61,62). Robust FISH staining was present in VA1-infected Caco-2 cells and was absent in mock-infected cells, validating our probe specificity (Supplemental figure 2). We next performed multiplexed FISH on heart tissue from VA1-infected and mock-infected WT A/J mice to determine the cellular tropisms. Each tissue was co-stained with the VA1 probe and with markers of cardiac myocytes (ryanodine receptor 2; Ryr2) and endothelial cells (von Willebrand Factor; vWF), and then imaged using confocal microscopy. We examined the tissue for cells co-expressing both VA1 RNA and cell-type markers in the same visual plane to determine the cell types that support viral replication. No signal was detected in the hearts of mock-infected mice, confirming the specificity of the VA1 probe in tissues (Figure 3B). Expected staining of the cardiac myocytes by Ryr2 and endothelial cells by vWF was detected (Figure 3B).  In VA1 inoculated mice, we detected intracellular viral RNA in cardiac cells, with the majority being myocytes as indicated by localization of Ryr2 and VA1 probes in the same cells (Figure 3C and 3D). Two types of staining patterns were observed for VA1. First, we noted punctate co-staining of both the VA1 and Ryr2 probes among clusters of densely packed nuclei without clear cellular borders (Figure 3C). This finding was consistent with the staining pattern from immunofluorescence to the viral capsid, which contained many nuclei based on DAPI staining (Figure 3A). These results suggest the possibility that increased nuclei is due to an inflammatory response that localizes to VA1-infected cardiomyocytes. In the second staining pattern, the probe localized to Ryr2 positive cells, suggestive of cardiac myocytes, without any apparent changes to cellular architecture with most of the cell positive for the VA1 probe (Figure 3D). These cells did not have any adjacent inflammatory infiltrates (Figure 3D).

Aside from myocytes, endothelial lineages of cells are common cardiac cell types (72). We also detected VA1 signal in regions that were expressing vWF and were negative for Ryr2 along the ventricular wall, suggestive of possible endocardial infection (Figure 3E). The positive VA1 staining was associated with infiltrating cells, consistent with the inflammatory foci that we hypothesized for the VA1-infected cardiac myocytes (Figures 3C and 3E). Together, these results clearly demonstrate that VA1 infects cells of the heart (Figures 3C-3E), is associated with increased nuclei suggesting an inflammatory response, and is not transiently passing through heart tissue or extracellularly attached.

In liver tissue, the *in situ* hybridization stains for viral RNA were negative (results not shown), suggesting a difference in the sensitivity of the *in situ* assay compared to qRT-PCR.

### Identification of histological features consistent with viral myocarditis

Given the detection of viral RNA in heart tissue and increased nuclei from immunofluorescence and FISH, we next determined if infection resulted in histological evidence of inflammation by examining hearts collected on day 7 and 21. Hematoxylin and eosin (H&E) stained sections showed focal interstitial clusters of lymphocytes within the myocardium (Figure 4A). The cellular infiltrates were predominantly mononuclear, lymphocytic, and were sporadically present, consistent with other animal models of viral myocarditis and in human myocarditis (Figure 4A) (69,73,74). Definitive cardiomyocyte damage was not observed in any cases. Analysis of the pericardium did not demonstrate any clear abnormalities. None of these findings were identified in mock-infected mouse tissue (Figure 4B). Analysis of heart tissue 21 days p.i. did not reveal any evidence of inflammatory infiltrates or other abnormalities of heart tissue (Figure 4C). When examining single tissue sections for each mouse, 2 of the 15 VA1-infected mice had foci of infiltrating immune cells at day 7 p.i., suggesting a mild myocarditis phenotype (Figure 4D).

**Fig 4.**
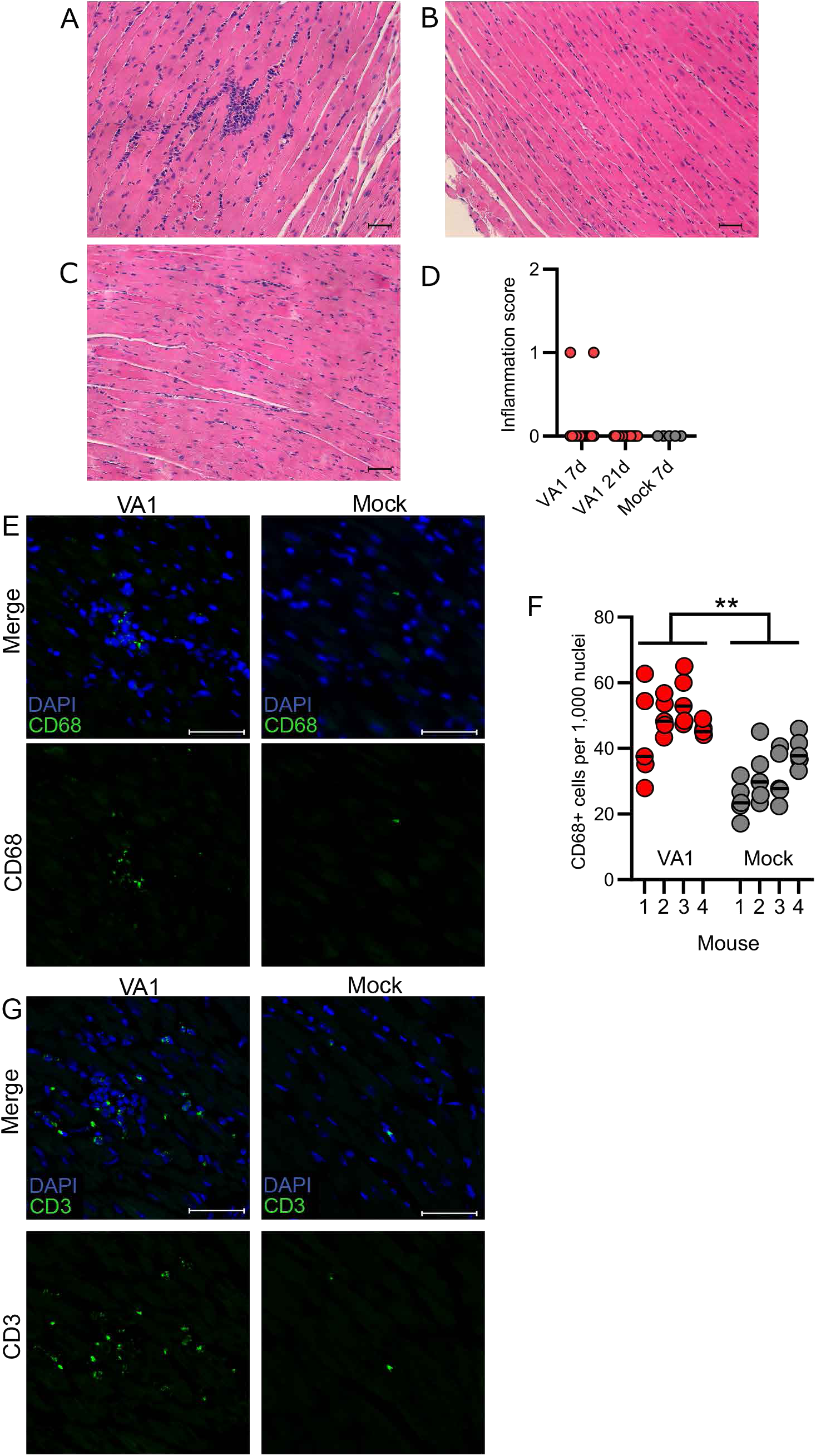
Histological findings consistent with myocarditis are present in VA1-infected heart tissue. Heart tissue histology was analyzed by hematoxylin and eosin staining with representative images shown. (A) Inflammatory infiltrates were present in some heart tissue on day 7, but (B) absent at day 21 or (C) in mock-inoculated mouse hearts. Scale bars represent 50 µm. (D) The presence of focal infiltrates was scored, and they were only present in hearts on day 7 post-inoculation. (E) Immunofluorescence was used to stain tissues for CD68, a marker of macrophages and other monocytes. Cells comprising the cellular infiltrates present in VA1-infected heart tissue were CD68 positive. Scale bars represent 20 µm. (F) CD68 positive cells were counted in representative tissue regions that lacked foci of cellular infiltrates. A significant increase in CD68 positive cells were identified in VA1-infected (mean 48.5 CD68+ cells per 1,000 nuclei) compared to mock infected hearts (mean 31.64 CD68+ cells per 1,000 nuclei; Nested T test t(6)= 4.35 P= 0.005). ** represents P ≤ 0.01. A total of four mice in each group were analyzed with 5 representative sections from each mouse used for counting per mouse. Horizontal lines represent median value. (G) Immunofluorescent staining for CD3, a marker of T cells. As with CD68, many of the cells with a focus of infiltrating cells were CD3 positive and were absent in mock infected mice. Scale bars are 20 µm.

We next stained the tissue for cellular markers at day 7 p.i. to determine the cellular types involved in the foci and whether other more subtle histological changes could be identified. We first stained the hearts using CD45 (PTPRC), a marker of nucleated hematopoietic cells (Supplemental Figure 3) (75). Robust staining of CD45 was detected from the cells in the foci of infiltrates, demonstrating their hematopoietic origin (Supplemental Figure 3). We next stained for macrophages (CD68) and T cells (CD3) as these cells are commonly involved in the pathogenesis of viral myocarditis and contribute to inflammatory infiltrates (76). When staining for CD68, we detected a proportion of cells in the foci were positive, demonstrating that some of the cells within the foci are macrophages (Figure 4E). In addition, we also noticed CD68 positive cells to be present diffusely through the heart tissue. When comparing VA1-infected to mock infected hearts in areas without infiltrating foci, there was an approximate 50% increase in CD68 positive cells (Nested T test t(6) = 4.35 P = 0.005; Figure 4F). We also detected that a significant proportion of cells in the foci were positive for CD3, consistent with lymphocytes as a cell population in the foci (Figure 4G). CD3 positive cells were not detected in significant quantity outside of the foci to allow for quantification. Taken together, these results demonstrate that macrophages and T cells are involved in the immune response to VA1 infection in the heart, consistent with other models of viral myocarditis (77). It also suggests that there is diffuse mobilization of macrophages into heart tissue, based on the increased number of CD68 positive cells.

Given the detectable viral loads in the liver, we also stained liver tissue on days 3 and 7 p.i.. The liver tissue did not demonstrate evidence of inflammation (data not shown).

### Immunodeficient mice are viremic and have increased viral RNA loads in heart tissue

Previous publications have demonstrated roles of the innate and adaptive immune responses during murine astrovirus infection (54,57,59). Recombinant activating gene 1 (Rag1) is a gene critical for recombination of B and T cell receptors, and knockout of Rag1 results in mice that are deficient in these cell lineages. Signal transducer and activation of transcription 1 (Stat1) knockout mice have a defect in interferon (IFN) response signaling, resulting in mice insensitive to IFN with reduced capacity to activate antiviral responses. Rag1 and Stat1 KO mice in the C57BL/6 background were inoculated with VA1 by IP inoculation, euthanized on day 7 p.i. and compared to WT C57BL/6 mice. Following inoculation, both Stat1 and Rag1 KO mice showed no clinical signs of infection or mortality. Both immunodeficient genotypes had 10-fold higher titers of virus in heart tissue (>10^5^ gRNA copies/mg) compared to WT (10^4^, Kruskal Wallis H(3)= 11.6, P = 0.003, post-hoc testing for Rag1 P = 0.006 and Stat1 P = 0.02; Figure 5A). In both immunodeficient backgrounds, viral capsid could be detected in heart tissue, with staining in cells consistent with cardiac myocytes (Figure 5B). Viral RNA was detected in cardiac myocytes as labeled by Ryr2 positive cells (example of Stat1 KO mouse; Figure 5C). We also detected in Rag1 KO mice that a subset of infected cells that were positive for vWF and negative for Ryr2 staining, consistent with possible endothelial infection (Figure 5D).

**Fig 5.**
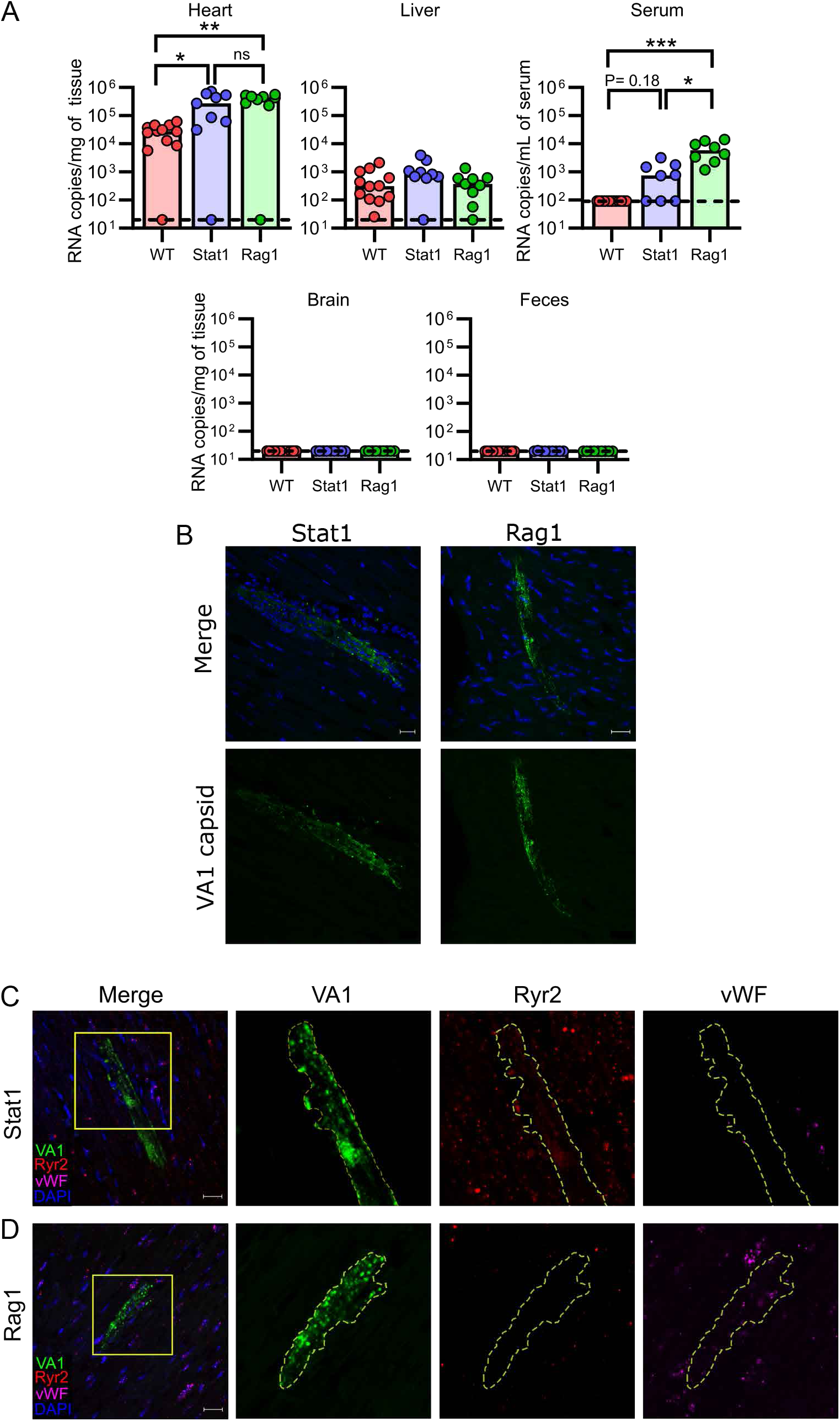
Immunodeficient mice support higher loads of VA1 in the heart and serum. Wildtype (WT) C57BL/6, Rag1 KO, and Stat1 KO mice were inoculated with VA1 by the IP route and tissues were collected 7 days post-inoculation. (A) VA1 copy number detected by qRT-PCR revealed higher quantities of VA1 RNA from heart tissue and serum of Rag1 KO mice compared to C57BL/6 WT (WT data from Fig 2). Stat1 KO mice had higher quantities of viral RNA in heart tissue. *** represents P value P value ≤ 0.001, ** represents P value ≤ 0.01, while * represents P value ≤ 0.05. (B) Immunofluorescence for VA1 capsid in immunodeficient mouse hearts. Both genetic backgrounds had positive staining for viral capsid, including large cell(s) with morphology consistent with cardiac myocytes. (C-D) Representative fluorescent *in situ* hybridizations for VA1 (green) in heart tissue, co-stained for cardiomyocytes (Ryr2 probe; red), and endothelial cells (vWF probe; magenta), with nuclei stained with DAPI (blue). Boxes highlight regions of interest that were further magnified. Scale bars represent 20 µm. Outlines highlight cells in which host markers localize to VA1 probes. (C) We detected VA1 RNA in cells expressing Ryr2, demonstrating infection cardiac myocytes in both Stat1 and Rag1 mice, with a representative infection depicted from a Stat1 mouse. (D) In Rag1 KO mice, we also identified VA1 infected cells (green) that were expressing vWF (magenta) but were not expressing Ryr2 (red), suggestive of endothelial cells.

Interestingly, immunodeficient mice were viremic (Kruskal Wallis H(3)= 20.5, P< 0.001; Figure 5A), with 100% of Rag1(post hoc testing P< 0.001) and ∼63% of Stat1 KO (post hoc testing P= 0.18) being viremic compared to none of the WT mice (Figure 5A). A statistical difference in the Stat1 KO compared to WT mice becomes apparent if binary positive/negative qRT-PCR results are used (Fisher’s Exact test P= 0.005). Both cohorts of immunodeficient mice had VA1 RNA present in the liver, while most other organs only had sporadic low-level detection of viral RNA, including the brain (Figure 5A), suggesting no additional tissue tropisms were identified compared to WT mice.

### Inflammatory foci in heart tissue are absent in Rag1 KO mice

Histopathological analysis of heart tissue from immunodeficient mice revealed differences between Stat1 and Rag1 KO mice. On H&E staining, Stat1 KO mice had multiple large cellular infiltrates while no infiltrates were identified in heart tissue from Rag1 KO mice (Figure 6A and 6B). We next stained the tissue for CD68 and CD3. The foci in Stat1 KO mice contained cells that were either CD68 or CD3 positive (Figure 6C and 6E). We also noted a significant increase in the number of CD68 positive cells compared to mock infection in fields without foci in the Stat1 KO mice (Nested T test t(6)= 5.65 P< 0.001; Figure 6D). In Rag1 KO mice, however, there was no increase in the number of CD68 positive cells compared to mock infection (Nested T test t(6)= 0.85 P= 0.43 ; Figure 6C and 6D). Not surprisingly, Rag1 KO mice had very rare staining for CD3, consistent with the rare viability of T cells from this mouse strain and expression of CD3 on other cell types including NKT cells (78,79). Collectively, these results indicate that both innate and adaptive immune systems are important in the antiviral response to VA1 infection. Infection of both immunodeficient mice result in increases in viral RNA, but only Stat1 KO mice can develop inflammatory foci and have increased number of CD68+ cells in heart tissue. The absence in Rag1 KO mice would suggest that lymphocytes are important for the formation of the foci and further recruitment of the monocytes/macrophages to heart tissue.

**Fig. 6.**
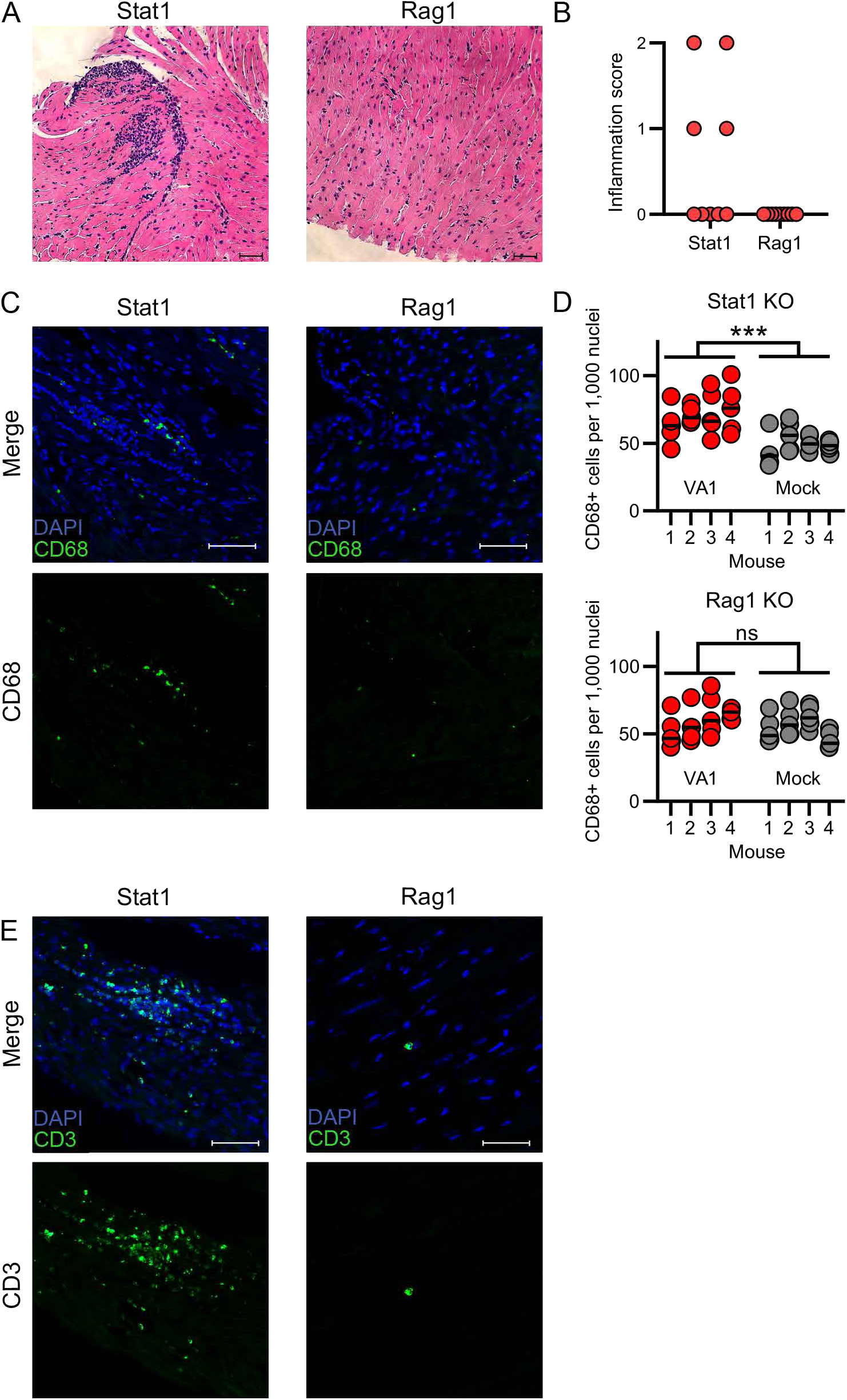
CD68 and CD3 positive cellular infiltrates are present in Stat1 KO but not Rag1 KO mice. (A-B) Hematoxylin and eosin staining of heart tissue collected from (A) Stat1 KO mice show clusters of cellular infiltration which are absent in Rag1 KO mice. Scale bars represent 50 µm. (B) Quantification of inflammation scores from Stat1 KO and Rag1 KO heart tissue 7 days post- inoculation. Foci were identified in Stat1 KO mice but none were present in Rag1 KO mice. (C) CD68 positive cellular infiltrates are present in Stat1 KO but not Rag1 KO mice as detected by immunofluorescence. Scale bars represent 50 µm (D) In representative tissue regions excluding foci of infiltrating cells, Stat1 (Nested T test t(6)= 5.65 P< 0.001) but not Rag1 KO mice (Nested T test t(6)= 0.85 P= 0.43) had significant increases in the number of CD68 positive cells compared to mock infected mice. Four mice per group were analyzed with 5 sections counted per mouse. Horizontal lines represent median value. (E) Immunofluorescence of heart tissue for CD3 with significant number of cells positive for CD3 in the infiltrating foci in Stat1 KO mice. Rare CD3 signal was detected in Rag1 KO mice. Scale bars represent 50 µm.

### VA1 can replicate in human cardiac endothelial cells *in vitro*

Based on the tropism we identified *in vivo*, we determined whether VA1 could replicate in primary murine or human cardiac myocytes (Figure 7A). In multi-step growth curves, there was no increase in VA1 RNA over time (Figure 7A). These results suggest that primary cardiac myocytes are non-permissive to infection *in vitro*, and additional factors affect the tropism we identified *in vivo*.

**Fig 7.**
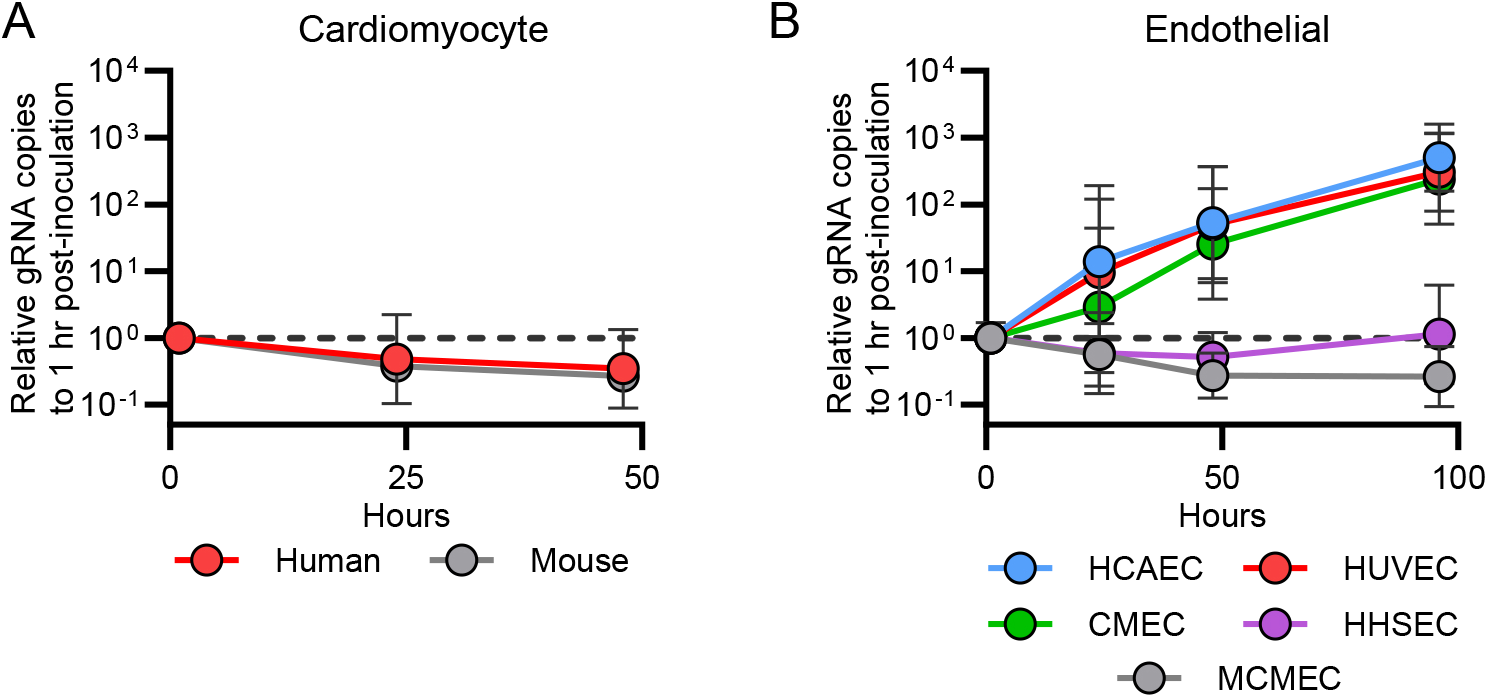
VA1 infects human endothelial-derived cell lines *in vitro*. Growth curves of VA1 using a multiplicity of infection of 3 in (A) primary human and mouse cardiac myocytes and (B) primary human endothelial cells including microvascular endothelial cells (CMEC), human coronary artery endothelial cells (HCAEC), human umbilical vein endothelial cells (HUVEC), and human hepatic sinusoidal endothelial cells (HHSEC), and mouse cardiac endothelial cells (MCMEC). Each data point is normalized to the gRNA copy number present at one hour post-inoculation for each cell line, and geometric means are plotted with error bars representing one geometric standard deviation. The horizontal dashed line represents the relative gRNA copy number at one hour post-inoculation.

The FISH assay also localized VA1 to the endocardium and endothelial cells. Endocardial cells are functionally related to endothelial cells (80). Based on the identification of an endothelial tropism, we tested the capacity of primary endothelial cells to support VA1 replication. We used primary human cardiac microvascular endothelial cells (CMEC), human coronary artery endothelial cells (HCAEC), and primary mouse cardiac microvascular endothelial cells (MCMEC).

We also tested two non-cardiac primary human endothelial cell lineages including umbilical vein (HUVEC) and hepatic sinusoidal cells (HHSEC). Interestingly, we identified a ∼100-fold increase in viral RNA in primary human CMEC, HCAEC, and HUVEC cells (Figure 7B). No significant replication was identified in HHSEC or MCMEC (Figure 7B). These results demonstrate that VA1 can replicate in human endothelial cells, including those derived from cardiac tissue, and further support the tropism identified from the mouse inoculations.

## Discussion

We describe here the first mouse model of infection using a human-infecting astrovirus species. Our data demonstrates that VA1 RNA accumulates within the hearts of infected mice, localizes intracellularly in myocytes, endothelial, and endocardial cells, resulting in translation of viral capsid, and can trigger an inflammatory response consistent with myocarditis. We also demonstrate reproducibility of this phenotype across multiple mouse genetic backgrounds, with greater viral loads in immunodeficient mice deficient in either innate or adaptive immunity. In addition, we showed that infectious virus could be recovered from heart tissue. Our results, through a combination of viral culture and molecular-based approaches, establish a causal relationship between astrovirus infection and cardiovascular disease in mice as they are consistent with the criteria established in Koch’s postulates and revised molecular-based guidelines (81,82).

These findings highlight the importance of animal models of infection, as this model revealed a novel cardiovascular tropism. Prior to this study, there was not a clear connection between astrovirus infection and cardiovascular disease. This has led to a paucity of testing and data regarding astrovirus infection of the heart. In a small number of studies, heart tissue from ducks, chickens, or geese infected with astroviruses were positive by PCR (35–37,39,83). In addition, human astrovirus was detected from heart tissue of one human case concerning for encephalitis (38). However, it is unknown if these results reflect transient presence of viral RNA in the tissue or blood, or *bona fide* infection because histological testing for virus was not completed. In Rawal *et al*, they reported findings from pigs with neurological disease associated with porcine astrovirus. In one animal, there was histological evidence of myocarditis, but PCR for astrovirus testing was negative from heart tissue (36). The cardiotropism of VA1 suggests that these previously reported findings may not be incidental and could be consistent with cardiovascular infection. Further epidemiological and experimental studies are necessary to determine whether the observed murine cardiotropism translates to cardiovascular diseases in humans and other organisms.Astroviruses could be a factor in cardiovascular health that has gone unrecognized.

Prior to this study, there also were major limitations for studying astroviruses *in vivo*. For the turkey astrovirus model, it is limited by the lack of turkey-specific reagents and clonal populations of turkeys (1). The mouse model of murine astrovirus is complicated by the frequent colonization of mice by murine astrovirus and the lack of a disease associated with infection (54,57,59). Previous attempts at generating an animal model for human astrovirus infection did not result in clear infection of the mice, with only transient detection of viral RNA in the spleen (54). Our model of VA1 infection demonstrates that human VA1 astrovirus can cause cross-species infection. This raises further hypotheses about conserved pathways between humans and mice that are utilized by the virus for the viral lifecycle, including whether the entry receptor(s) for VA1 are the same or different between species. It is also not clear whether the myocarditis phenotype is specific to VA1 or if other human-infecting astroviruses cause myocarditis in mice. It is also unclear if VA1 could be used for inoculation of well-studied organisms, including non-human primates, and exhibit the same cardiotropism.

This study provides important immunological insights into the response to human astrovirus infection. We identified increased viral RNA loads in heart tissue and viremia from Rag1 KO mice, supporting the role of the adaptive immune response during infection. We also did not identify any inflammatory foci or increase in CD68+ cells in the Rag1 KO mice. This suggests that formation of the foci and further recruitment of monocytes/macrophages during VA1 infection is mediated by lymphocytes. Lymphocyte-mediated elimination of virally infected cells could be a mechanism that explains the increase in viral RNA. Increased viral loads have been also identified in models of murine astrovirus infection using mice with genetic defects in the adaptive immune response (54,57). In humans, a majority of the patients with astrovirus-associated encephalitis had defects of adaptive immunity. Some subjects had a primary immunodeficiency including X-linked agammaglobulinemia, while others had acquired immunodeficiencies due to recently receiving chemotherapy or a hematopoietic stem cell transplant (1). Astroviruses could be opportunistic pathogens in humans. The lack of a coordinated lymphocyte response could lead to increased viral replication and viremia, thus promoting invasive diseases. Components of cellular immunity, including cytotoxicity and downstream processes to clear virus enabled by effector T cells, could be important for the immune response and can be further studied. Future work will also determine the role of antibody-mediated neutralization and Fc effector functions.

Our results also suggest the importance for the innate immune response to VA1 infection as we observed higher quantities of VA1 RNA in Stat1 KO mice compared to WT mice. This result is consistent with past results where we have previously shown *in vitro* that VA1 is IFN sensitive (61,84). IFN is also important in the models of murine astrovirus infection, where higher quantities of murine astrovirus RNA was detected in IFN-deficient mouse backgrounds (54,57). We hypothesize that the increase in viral RNA in Stat1 KO mice is due to disruption of IFN signaling leading to a diminished innate immune response (85). However, we cannot rule out an alternative possibility that Stat1 could be mediating effects through adaptive immunity, as Stat1 also has a role in antigen presentation and antibody class switching (86). Nonetheless, the Stat1 KO mice had inflammatory foci containing T cells and increased numbers of macrophages that were also present in the WT mice. These results suggest the Stat1 KO mice are mounting an adaptive immune response to some extent. Future work will dissect the mechanisms behind why Stat1 KO mice have an increase in VA1 RNA, identifying whether it is due to a reduced innate immune response or is mediated by an alternative pathway.

The VA1 myocarditis model revealed novel tissue tropisms of VA1 infection in cardiomyocytes, endothelial, and endocardial cells. This provides a direct route by which VA1 could cause cardiac injury by mediating an inflammatory response. In addition, VA1 replicates in multiple human endothelial cell lines. The endothelial tropism has many pathological consequences as other viral infections of endothelial cells promote expression of pro-inflammatory cytokines and chemokines (87,88) that can be now further evaluated *in vitro* and *vivo*. Inoculation of VA1 into primary cultures of murine myocytes and endothelial cells did not support viral replication. Viral *in vivo* tropisms do not always directly translate to *in vitro* tropisms, as decades of work were needed to propagate human norovirus in GI cells and hepatitis C in hepatic cells (89,90). Furthermore, murine astrovirus has been difficult to cultivate *in vitro* despite clear evidence of infection of the gastrointestinal tract *in vivo* (91,92). Murine astrovirus is incapable of propagation in three dimensional enteroids and there is only around a 10-fold increase in viral RNA using air-liquid interface cultures derived from mouse enteroids (91). There are likely additional factors yet to be identified that contribute to the tropisms of astroviruses *in vitro* and *in vivo* that affect their capacity to replicate in these different models.

Despite the known fecal-oral transmission route and association of gastroenteritis with astroviruses in humans, mice are largely resistant to PO and OG challenge with VA1. A similar result has been published with other human-infecting astroviruses (54). While VA1 infected the heart in IP inoculated mice, we did not observe significant infection of the GI tract. The factors conferring the resistance of the GI tract are unknown. Mice can be resistant to oral challenge but susceptible to other routes of inoculation with other well-known fecal-orally transmitted viruses in humans, including many enteroviruses (93). Currently the receptor for VA1 is unknown, so it is unclear whether the entry receptor affects the tropism to the GI tract, and therefore susceptibility. In addition, Ingle *et al* previously demonstrated that laboratory mice were resistant to infection with murine norovirus when the mice were co-infected with a specific murine astrovirus strain (94). Murine astrovirus infection was associated with elevated IFN-λ, which was hypothesized to prevent infection with norovirus or other viruses (94). Interestingly, in our IP infections of Stat1 KO mice (Figure 5A), we did not detect any VA1 RNA in the feces. Stat1 is important for the signaling cascade for IFN-λ (95), so these results would not be consistent with the hypothesis that IFN-λ is restricting VA1 infection of the GI tract. In the future, we can further determine how host factors like the entry receptor, IFN-λ, and other confounders like murine astrovirus infection affect the infectability and tropism of VA1 *in vivo*.

The VA1 mouse model does have limitations in modeling astrovirus associated-human diseases. Astroviruses causes encephalitis in humans, but we note the lack of clear evidence of VA1 infection in the central nervous system by either IP or IC routes of inoculation. Other human neurotropic viruses have shown to lack a neurotropism in mice or required adaptation to cause neurological disease including poliovirus, measles, enterovirus 71, and enterovirus D68 (96–100). We have previously shown in cell culture that VA1 can infect astrocytes, but neurons are resistant to infection, despite a neuronal tropism being identified from clinical cases (11,62). In addition, VA1 did not cause significant infection of the GI tract despite the link of astroviruses with gastrointestinal diseases (3). Together, these results suggest there are additional unknown factors that affect the viral tropism *in vitro* and *in vivo*, and they can be further characterized in the future. In addition, it is possible that further adaptation of the virus to mice is necessary to observe these diseases. Future work can focus on serially passaging VA1 in mice to determine if the virus accumulates adaptive mutations that confer a change in tropism or pathogenicity in mice. Astrovirus infection in humans is very common based on seroprevalence studies. We have established a tractable mouse model of VA1 infection that will be an important tool to study the pathogenesis of human astrovirus infection *in vivo.* In addition, the model also establishes a previously unappreciated cardiovascular tropism in mice, creating additional hypotheses regarding the role of astroviruses in cardiovascular health. Ultimately, this model enables further understanding of the biology of astroviruses *in vivo* and will allow for future studies in the development of astrovirus-specific therapeutics and vaccines.

## Funding

This work was supported by the following: A. B. J. received support from National Institute of Allergy and Infectious Disease [K08 AI132745], by the National Center for Advancing Translational Sciences of the National Institutes of Health under Award Number [UL1 TR002345], and the Children’s Discovery Institute of Washington University and St. Louis Children’s Hospital. The funders had no role in study design, data collection and analysis, decision to publish, or preparation of the manuscript.

## Acknowledgements

We would like to thank the various contributions of Herbert “Skip” Virgin over the years, including his technical advice and support in establishing the mouse inoculation protocols.

## Declaration of interest

none

---

**Supplemental Fig 1.**
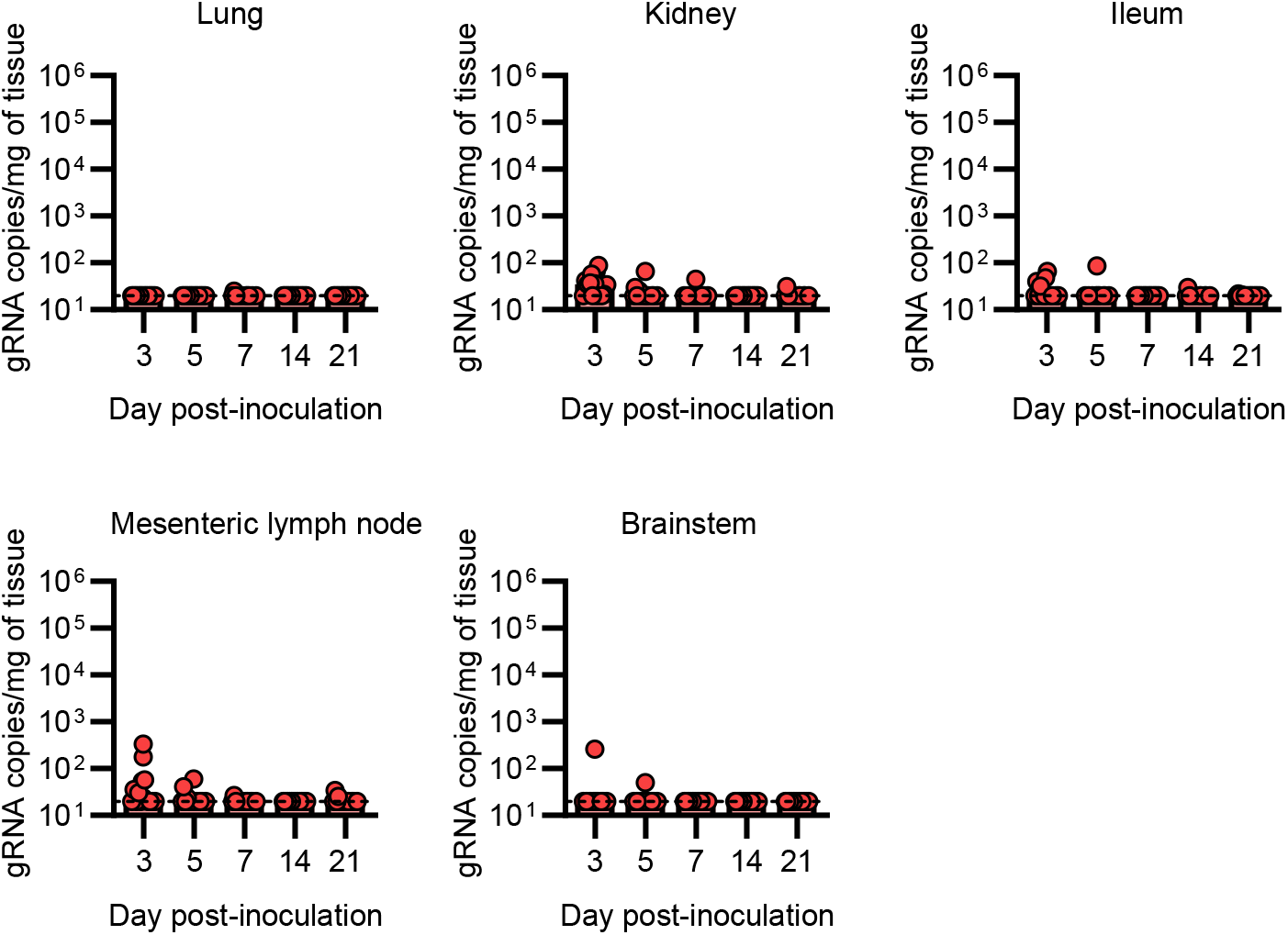
VA1 is sporadically detected in low quantities in various tissues over a 21 day time course following inoculation Viral RNA copy numbers were measured by qRT-PCR from multiple additional tissues after IP inoculation of VA1 in wildtype A/J mice. Most tissues were undetectable at any time point, including the lung, kidney, ileum, mesenteric lymph node, and brainstem tissue. Dashed line represents the limit of detection.

**Supplemental Fig 2.**
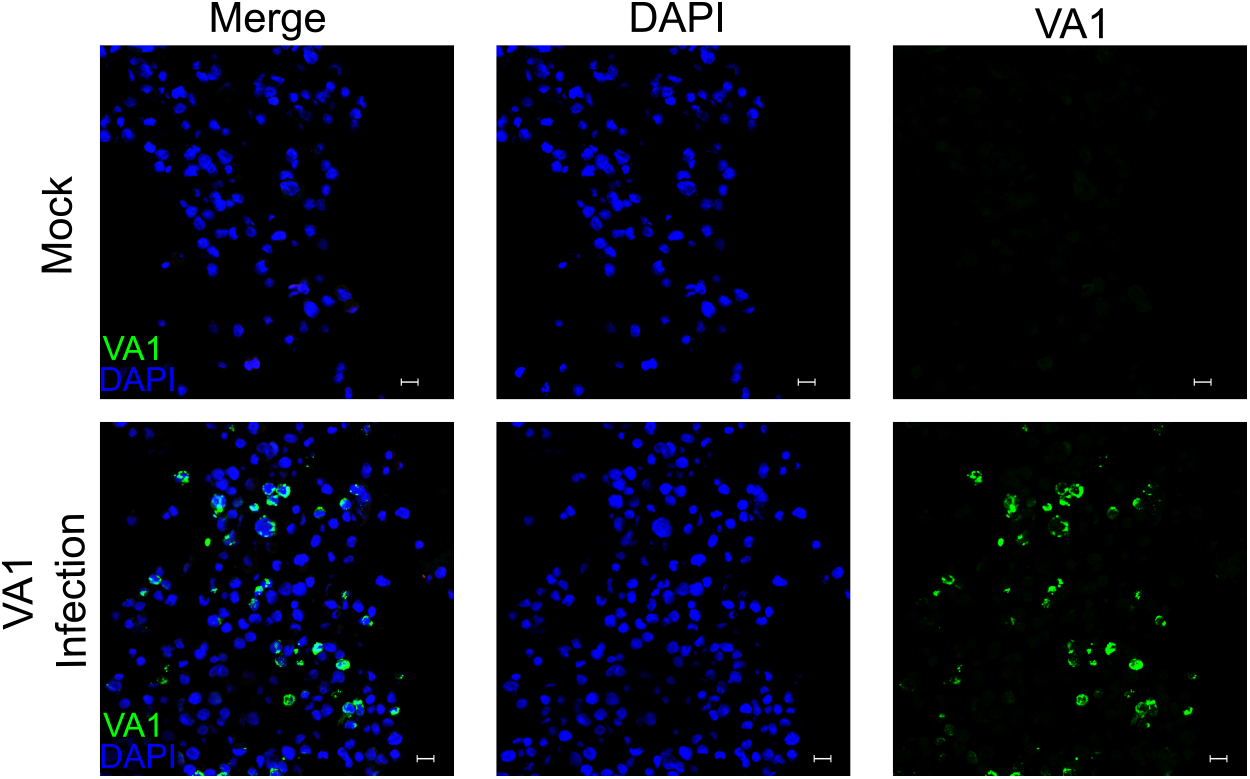
Development of a fluorescent *in situ* hybridization (FISH) assay A fluorescent in situ hybridization (FISH) assay was developed using probes complementary to ORF2 of VA1. VA1-infected or mock-infected Caco-2 cells were stained with the VA1 probe and DAPI (blue), demonstrating specificity of the probes to virally infected cells. Scale bars represent 20 µm.

**Supplemental Fig 3.**
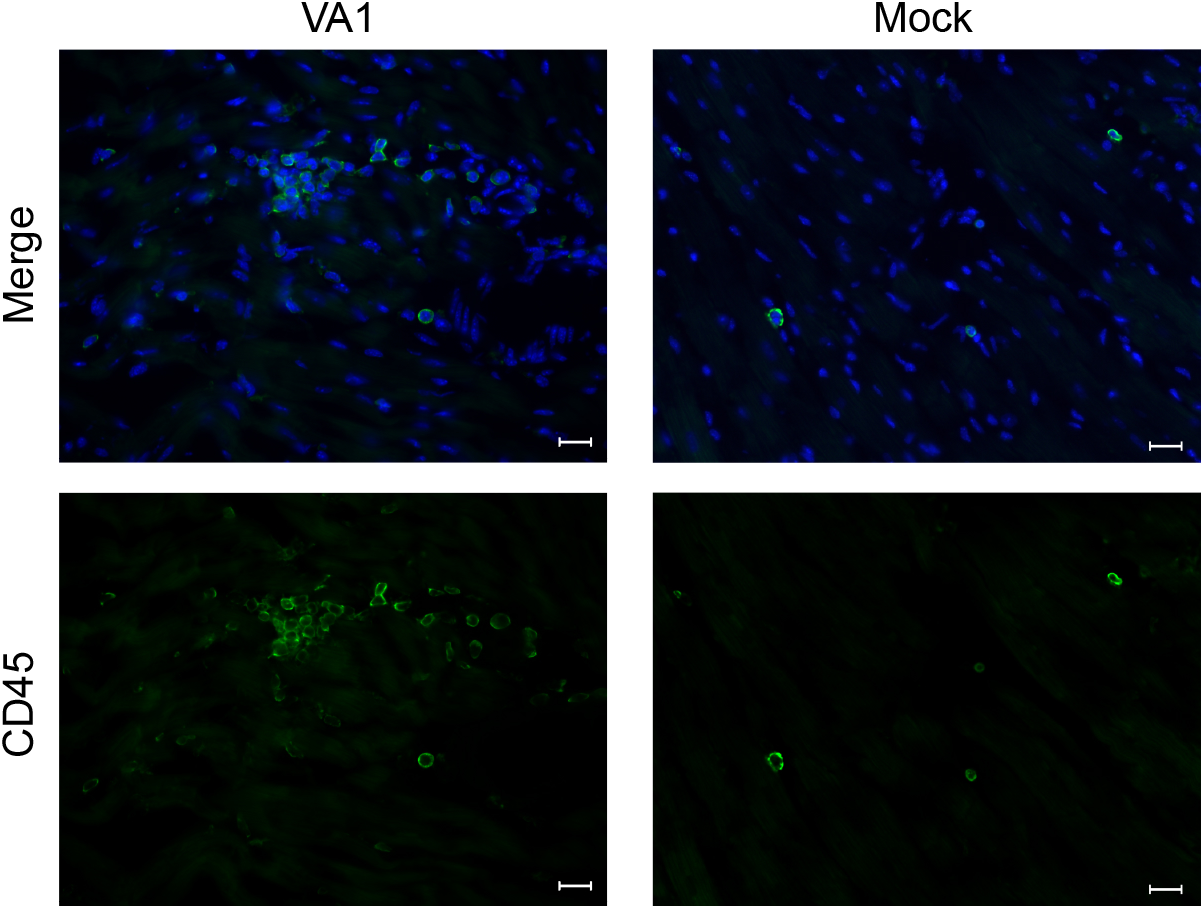
Cellular infiltrates in heart tissue are CD45 positive Heart tissue sections from VA1 or mock-infected mice were stained for CD45. Foci of infiltrating cells were only identified in VA1 infected heart tissue and stained positive for CD45. Only sporadic detection of CD45 was detected in mock-infected heart tissue. 40x magnification, scale bars represent 20 µm.

## Notes

### Competing Interest Statement

The authors have declared no competing interest.

